# Characterization of histone inheritance patterns in the *Drosophila* female germline

**DOI:** 10.1101/2020.08.18.255455

**Authors:** Elizabeth W. Kahney, Lydia Sohn, Kayla Viets-Layng, Robert Johnston, Xin Chen

## Abstract

Stem cells have the unique ability to undergo asymmetric division which produces two daughter cells that are genetically identical, but commit to different cell fates. The loss of this balanced asymmetric outcome can lead to many diseases, including cancer and tissue dystrophy. Understanding this tightly regulated process is crucial in developing methods to treat these abnormalities. Here, we report that produced from a *Drosophila* female germline stem cell asymmetric division, the two daughter cells differentially inherit histones at key genes related to either maintaining the stem cell state or promoting differentiation, but not at constitutively active or silenced genes. We combined histone labeling with DNA Oligopaints to distinguish old versus new histone distribution and visualize their inheritance patterns at single-gene resolution in asymmetrically dividing cells *in vivo*. This strategy can be widely applied to other biological contexts involving cell fate establishment during development or tissue homeostasis in multicellular organisms.

## INTRODUCTION

Stem cells often undergo asymmetric cell division to give rise to a daughter that self-renews and another daughter that differentiates, despite each containing identical genomes. The differences in cell fate can be achieved through epigenetic mechanisms, defined as modifications that change gene expression without altering the primary DNA sequences. Epigenetic modifications can be maintained even after going through multiple cell divisions, a phenomenon called “cellular memory” (Jacobs & van Lohuizen, 2002, Probst, Dunleavy et al., 2009, Ringrose & Paro, 2004, Turner, 2002). Mis-regulation of this process can be detrimental, leading to many diseases including cancers, tissue dystrophy, and infertility (Baylin & Ohm, 2006, Chambers, Shaw et al., 2007, Feinberg, Ohlsson et al., 2006, Fitzsimons, van Bodegraven et al., 2014, Fredly, Gjertsen et al., 2013). However, the processes by which epigenetic information is inherited or changed during cell divisions in endogenous stem cells or asymmetrically dividing cells are still not well understood.

The known primary mechanisms of epigenetic regulation include DNA methylation, chromatin remodeling, post-translational modifications of histones, histone variants, and non-coding RNAs (Jin, Li et al., 2011, Kouzarides, 2007, Lee, 2012). Since only about 0.07% of the genomic DNA in *Drosophila* is methylated in adult flies, it is likely that histones carry the majority of epigenetic information in the adult germline (Lyko, Ramsahoye et al., 2000, Zhang, Huang et al., 2015). When the (H3-H4)2 tetramer combines with two H2A/H2B dimers, an octamer encircled by DNA structure forms, which is called nucleosome, the basic unit of chromatin structure. The N-terminal tails of histones can undergo extensive post-translational modifications that can change the local structure of chromatin and the expression of the genes therein. Notably, most post-translational histone modifications that regulate gene expression are found on the N-termini of H3 and H4 (Allis & Jenuwein, 2016, Huang, Sabari et al., 2014, Janssen, Sidoli et al., 2017, Kouzarides, 2007, Young, DiMaggio et al., 2010). Furthermore, the age of a histone protein determines the specific post-translational modifications it carries (Alabert, Barth et al., 2015, Xu, Wang et al., 2011, Zee, Britton et al., 2012). The molecular mechanisms by which newly synthesized (new) histones are incorporated onto DNA has been well studied, but little is known about how pre-existing (old) histones are re-incorporated onto DNA following replication and subsequently segregated during cellular division (Ahmad & Henikoff, 2018, Alabert & Groth, 2012, Serra-Cardona & Zhang, 2018). Recent studies have shed light on both the molecular mechanisms (Gan, Serra-Cardona et al., 2018, Petryk, Dalby et al., 2018, Yu, Gan et al., 2018) and cellular responses (Escobar, Oksuz et al., 2019, Lin, Yuan et al., 2016, Reveron-Gomez, Gonzalez-Aguilera et al., 2018) responsible for recycling old histones in symmetrically dividing cells. However, it remains elusive how old versus new histones are partitioned in asymmetrically dividing cells, such as certain types of adult stem cells. For example, stem cells may maintain their identity by selectively inheriting particular post-translational modifications on old histones to sustain an active transcription at stemness-promoting genes but a repressive state at differentiation and non-lineage-specific genes. On the other hand, differentiating daughter cells may inherit new histones to reset the chromatin in preparation for cellular differentiation.

The *Drosophila* germline stem cell (GSC) systems permits the visualization of asymmetric cell division (ACD) at a single-cell resolution *in vivo* (Fuller & Spradling, 2007). In both male and female gonads, GSCs can divide asymmetrically to produce a self-renewed GSC and a cystoblast (CB, in the female germline), or a gonialblast (GB, in the male germline), that subsequently undergoes a transit-amplification stage of four mitoses before entering meiosis and terminal differentiation. During ACD in *Drosophila* male GSCs, the old histone H3 is selectively segregated to the renewed stem cell, whereas the new H3 is enriched in the GB (Tran, Lim et al., 2012). Mis-regulation of this differential inheritance can lead to stem cell loss and early-stage germline tumors (Xie, Wooten et al., 2015). The asymmetric inheritance of old versus new H3 is specific to asymmetrically dividing GSCs and occurs with histones H3 and H4, but not with H2A, and such patterns are established during DNA replication (Wooten, Snedeker et al., 2019). Furthermore, biased sister chromatid segregation in male GSCs are regulated by the coordination between dynamic microtubule activity and asymmetric sister centromeres (Ranjan, Snedeker et al., 2019). It is possible that the asymmetric inheritance of old (H3-H4)2 tetramers versus new (H3-H4)2 tetramers could be responsible for differential gene expression in genetically identical daughter cells.

However, the question remains whether differential histone inheritance is a conserved feature of stem cells and/or asymmetrically dividing cells or not. The *Drosophila* female germline is an excellent system to address this question. *Drosophila* ovaries consist of 16-18 ovarioles, each of which contain an anatomically defined stem cell niche that supports 2-3 GSCs, which can undergo ACD at their apical tips (Xie & Spradling, 2000). Both *Drosophila* male and female germlines have well-characterized cellular features. Recent studies revealed that asymmetric microtubules closely interact with asymmetric sister centromeres to establish biased attachment and the segregation of sister chromatids in female GSCs (Dattoli, 2020), resembling the results in male GSCs (Ranjan et al., 2019). However, there are key differences between the male and female systems. For example, in males, one GB divides four times to produce 16 spermatogonial cells, all of which subsequently undergo meiosis to produce a total of 64 sperm (Hardy, Tokuyasu et al., 1981, Hardy, Tokuyasu et al., 1979, Tokuyasu, Peacock et al., 1977).

By contrast, the CB in females undergoes four rounds of mitosis but produces two pre-oocytes and 14 nurse cells. One of the pre-oocytes is determined as the oocyte and undergoes meiosis, whereas the other pre-oocyte joins the nurse cell fate. Together, all 15 nurse cells undergo endoreplication and produce mRNAs and proteins to support oogenesis and early embryogenesis (King, 1957, Smith & Orr-Weaver, 1991). Interestingly, the initial division of the CB produces the two pre-oocytes, but only one eventually takes on the oocyte fate. It has been suggested that the CB division is asymmetric, which would create oocyte versus nurse cell fate, and that spectrosome inheritance may play a role in such a cell fate determination (de Cuevas & Spradling, 1998).

Here, we show that in both *Drosophila* female GSC and CB divisions, large chromosomal regions carrying distinct old versus new histone H3 can be detected. This feature gradually disappears throughout subsequent mitotic divisions with higher degree of cellular differentiation, giving support to the hypothesis that both the GSC and CB division may involve the differential partitioning of intrinsic factors such as histone information, which may contribute to the differences in their daughter cell fates. Additionally, we combined DNA Oligopaints with a two-color histone labeling system to visualize genomic region-specific histone inheritance patterns in mitotically dividing female germ cells *in vivo*. We found that the two daughter cells produced from a GSC division differentially inherit histones at genomic regions harboring key genes, such as *daughters against dpp* (*dad*) for maintaining the stem cell state (Casanueva & Ferguson, 2004, Kirilly, Wang et al., 2011, Xie & Spradling, 1998) and *bag of marbles* (*bam*) for promoting differentiation (McKearin & Ohlstein, 1995, McKearin & Spradling, 1990). Overall, this study offers insight into how epigenetic information is inherited through the process of asymmetric cell division and offers a technique to simultaneously visualize genetic and epigenetic information. The techniques used here are applicable to many other stem cell lineages and organisms, which can provide valuable insight into the molecular mechanisms and biological significance of ACD.

## RESULTS AND DISCUSSION

### Old versus new histone H3 display non-overlapping patterns in dividing female GSCs, but not in dividing cystocytes

The *Drosophila* female germline is an excellent system to study germ cell differentiation at a single-cell resolution (Fig 1A). To investigate histone inheritance patterns in early-stage female germ cells including GSCs, we used a heat-shock controlled, dual-fluorescent, tag-switching histone transgene driven by *GreenEye-nanos-Gal4* which drives transgene expression strongly in the early germline of both male and female *Drosophila melanogaster* [This driver is named such because it was originally paired with a synthetic *eyeless* promoter, 3xP3>eGFP, as a fluorescent reporter (Holtzman, Miller et al., 2010)]. Using this labeling system as described previously (Ranjan et al., 2019, Wooten et al., 2019), we tagged old histones with eGFP (green fluorescent protein) and new histones with mCherry (red fluorescent protein). The switch from eGFP to mCherry labeling is controlled by a FLP recombinase, driven by a heat shock promoter [Fig 1B, (Tran et al., 2012)]. The cell cycle of female GSCs is approximately 24 hours without synchrony, with G2 being the longest phase, lasting for 18-20 hours (Ables & Drummond-Barbosa, 2013, Hinnant, Alvarez et al., 2017, Hsu, LaFever et al., 2008). Therefore, it is likely that the heat-shock induced histone tag switch occurs in the G2 phase. Allowing a recovery time of 34-40 hours post heat-shock ensures that the tag-switched GSCs finish another complete round of DNA replication, where mCherry-labeled new histones are incorporated genome-wide. GSCs then advance into the subsequent mitosis, at which point the distribution of old versus new histones in female GSCs is examined (Fig 1C).

**Figure 1:**
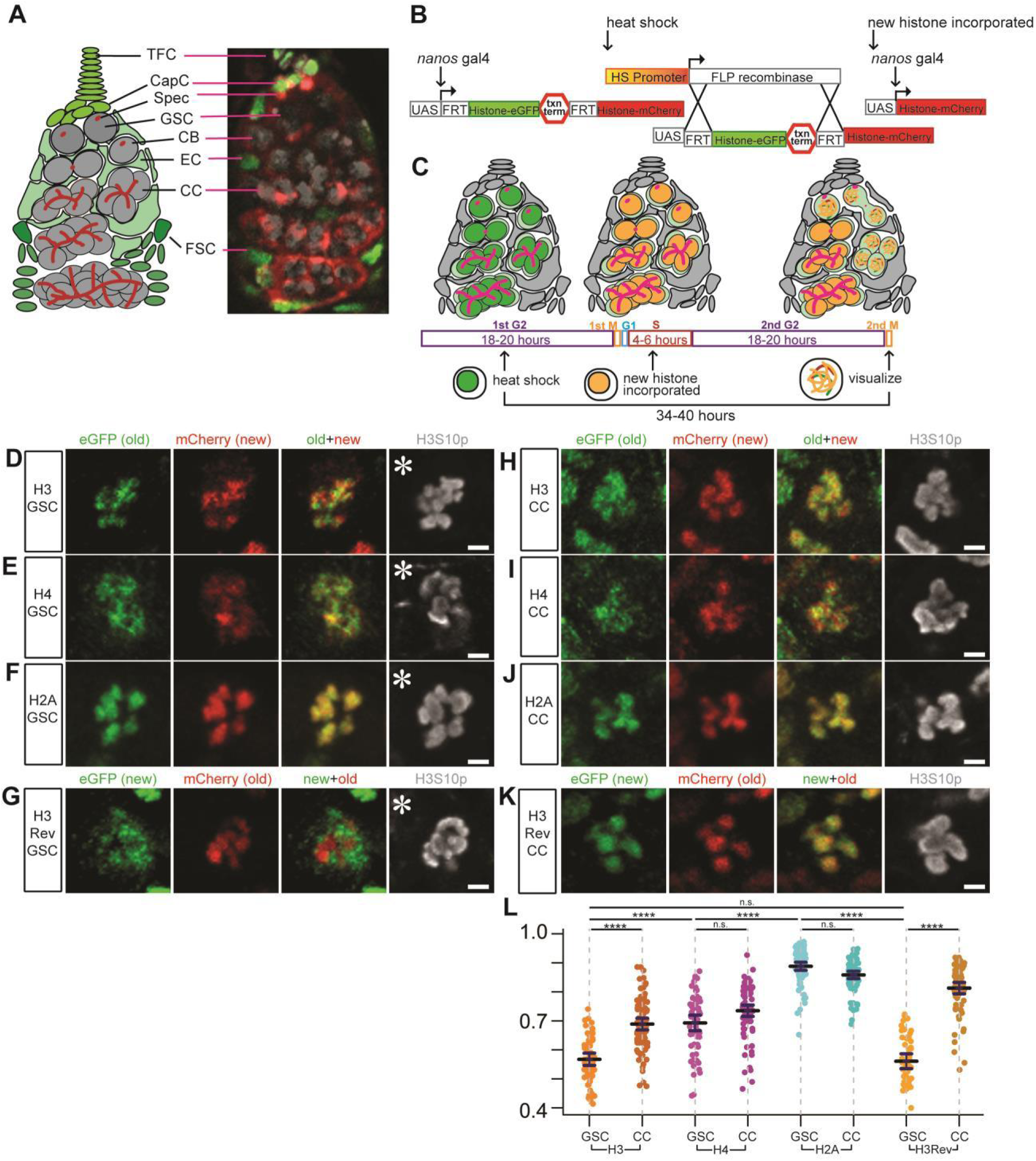
Non-overlapping old versus new histone H3 distribution patterns in mitotic *Drosophila* female germline stem cells (GSCs), but not in cystocytes (CCs). (**A**) A cartoon and corresponding immunofluorescence image depicting the *Drosophila* germarium. Terminal filament cells (TFC) and cap cells (CapC) create a niche for female germline stem cells (GSC), which divide asymmetrically to self-renew and produce a cystoblast (CB). The CB undergoes four mitotic divisions to create differentiating cystocytes (CC). The germline (gray) is supported by somatic (green) escort cells (EC) and follicle stem cells (FSC) that produce follicle cells that surround developing egg chambers. The spectrosome (Spec, red) is a specialized organelle in the early-stage germline, such as GSCs and CBs. The spectrosome structure is round while fusome is branched, which branches over the mitotic divisions in further differentiated germ cells, connecting the CCs within a cyst. (**B**) A cartoon detailing how the heat shock controlled dual-color system driven by *GreenEye-nanos-Gal4* labels preexisting (old) histones with eGFP (green fluorescent protein) and newly synthesized (new) histones with mCherry (red fluorescent protein), respectively. (**C**) A scheme of recovery time and histone incorporation after heat shock that induces an irreversible genetic switch in the histone transgene. (**D-F**) Old (green) versus new (red) histone patterns for H3 (**D**), H4 (**E**), and H2A (**F**) in prometaphase female GSCs marked by anti-H3S10ph (gray). (**G**) Old (red) versus new (green) histone patterns for H3Rev in prometaphase female GSCs marked by anti-H3S10ph (gray). (**H-J**) Old (green) versus new (red) histone patterns for H3 (**H**), H4 (**I**), and H2A (**J**) in prometaphase female 4-cell CC marked by anti-H3S10ph (gray). (**K**) Old (red) versus new (green) histone patterns for H3Rev in prometaphase female 4-cell CC marked by anti-H3S10ph (gray). (**L**) Quantification of the overlap degree between old and new histones in late prophase and prometaphase GSCs and CCs using Spearman correlation: The measurement is from a single Z-slice at the center of each mitotic nucleus, which shows results similar to analyzing every Z-slice throughout the entire Z-stacks followed by averaging them (Fig EV1A, see Materials and Methods). Values are mean ± 95% CI. *P*-value: pairwise ANOVA test with bonferroni correction. ****: *P*< 0.0001, n.s.: not significant. Asterisk: niche. Scale bars: 2μm. See relevant Supplemental Table for individual data points for **Fig 1L**.

Using a mitotic marker (anti-H3S10phos or H3S10ph) and chromosomal morphology to identify late prophase and prometaphase GSCs, we found that old and new H3 occupied distinct chromosomal domains shown as separable eGFP (old H3) and mCherry (new H3) signals on condensed chromosomes (old + new in Fig 1D). This separation was also detected in a control H3 line in which the old and new fluorescent tags were reversed (H3Rev: mCherry labeled old H3 and eGFP labeled new H3, Fig 1G). Contrastingly, more overlapping signals were detected in mitotic GSCs where old histone H2A was labeled with eGFP and new H2A by mCherry (Fig 1F). Finally, mitotic GSCs expressing old H4 (eGFP) and new H4 (mCherry) displayed a moderate separation (Fig 1E): Old and new H4 had less overlap than old and new H2A (Fig 1F) but more overlap than old and new H3 (Fig 1D).

In order to quantify these imaging results, we measured Spearman’s rank correlation coefficient (Schober, Boer et al., 2018) for old versus new H3, H4, H2A, and H3Rev, respectively (Fig 1L). Old and new H3 consistently showed the lowest correlation coefficients of 0.567 (*n*= 57) in the H3 line and of 0.561 (*n*= 50) in the H3Rev line, indicative of the highest separation between these two signals. By contrast, old and new H2A displayed the highest correlation coefficient of 0.888 (*n*=79), suggesting the highest overlap between these two signals. Consistent with the moderate separation between old and new H4, the average correlation coefficient of 0.693 (*n*= 60) is between that of H3 and H2A. Although canonical H3 and H4 form tetramers *in vivo*, their different behaviors could be due to H4 pairing with both H3 and the histone variant H3.3.

To directly test this possibility, we examined the correlation coefficient of old versus new H3.3 using a similar labeling scheme and parallel analysis. Since new H3.3 is incorporated in a replication-independent manner (Ahmad & Henikoff, 2002a, Ahmad & Henikoff, 2002b, Tagami, Ray-Gallet et al., 2004), we recovered female flies for 18-20 hours (∼ one cell cycle) after the heat shock-induced *H3*.*3-eGFP* to *H3*.*3-mCherry* expression switch and subsequently examined the mitotic GSCs. Indeed, an average correlation coefficient of 0.752 was detected for old and new H3.3 (*n*=55, Fig EV1E). The correlation coefficient of H4 in mitotic GSCs (0.693) is very close to the average of the H3 and H3.3 correlation coefficients at 0.660. This supports the hypothesis that the moderate separation between old and new H4 is due to the different overlapping degrees of its binding partners: low for old versus new H3 and high for old versus new H3.3.

Next, to determine whether the non-overlapping old versus new H3 distribution patterns have cellular specificity, we imaged dividing cystocytes (CCs) and quantified them at the 4-cell and 8-cell stage (Fig 1L). For H3, late prophase to prometaphase CCs showed an average Spearman correlation coefficient of 0.689 (*n*=83), indicating a significantly higher degree of overlap (or colocalization) between old and new H3 in CCs than in GSCs (Fig 1H versus Fig 1D, quantified in Fig 1L). The H3Rev also had a significantly higher correlation coefficient of 0.813 (*n*=76) in late prophase to prometaphase CCs compared to GSCs (Fig 1K versus Fig 1G, quantified in Fig 1L). Old and new H4 displayed an average correlation coefficient of 0.735 (*n*=81) in mitotic CCs, which was not significantly higher than the correlation coefficient in mitotic GSCs (Fig 1I versus Fig 1E, quantified in Fig 1L). Lastly, the degree of overlap between old and new H2A was indistinguishable between GSCs and CCs, with the latter having an average correlation coefficient of 0.858 (*n*=79, Fig 1J versus Fig 1F, quantified in Fig 1L).

Taken together, these data demonstrate that the non-overlapping old versus new histone domains seen for H3 are more prominent in GSCs than in CCs. The cellular specificity of the non-overlapping old versus new histone H3 patterns recapitulates what has been previously reported in *Drosophila* male GSCs, where the global asymmetric inheritance of old versus new H3 is specifically found in asymmetrically dividing GSCs but not in symmetrically dividing spermatogonial cells (Tran et al., 2012).

### Old versus new histone distribution patterns during mitosis reveal CBs are more like GSCs than CCs

Given that old and new H3 histones appear in separable domains in GSCs but not in CCs (Fig 1), we further examined the histone distribution patterns for H3, H4, H2A, H3.3, and H3Rev in mitotic cells at each stage during transit amplification in the female germline lineage and compared them with the distribution patterns in GSCs. Interestingly, we found that the separable old versus new H3 pattern gradually diminishes as germ cells differentiate (Fig EV1B for H3, EV1C for H3Rev). Specifically, the old versus new H3 correlation coefficient does not significantly differ between GSCs and CBs in both H3 lines, suggesting that this separation is independent of the tag but dependent on the different behaviors between old and new H3 proteins in early-stage female germ cells. Additionally, both lines display a significantly increased correlation coefficient from the early staged GSC and CB cells to the 2-cell stage and subsequently to the 4-cell stage, suggesting more overlap between old and new H3 as germ cells differentiate. Furthermore, the gradual change of the old versus new histone correlation coefficient during germline differentiation is specific to the histone H3 (Fig EV1B-C). Other histones, including H4, H3.3, and H2A (Fig EV1D-F), showed no significant differences in the overlap of old versus new histone protein between each consecutive cell stage, although there is a statistically significant decrease between H2A in GSCs and CBs (Fig EV1F, see Figure legend for possible reasons). However, the correlation coefficients for H2A in GSCs and CBs were both significantly higher than the correlation coefficients for H3 in the same staged cells (Fig EV1G).

Previously, it has been proposed that the CB division could be a “symmetry breaking” step, as only one of the two daughter cells from this division eventually differentiates into the oocyte. Therefore, it is possible that breaking the symmetry in preparation for cellular differentiation in the female germline lineage is accomplished by two steps, GSC division and CB division. To further explore this possibility, we next studied old versus new H3 segregation patterns during anaphase and telophase in GSCs and CBs.

### Dividing female GSCs and CBs show globally symmetric but locally asymmetric segregation patterns of old versus new histone

Using the H3S10ph mitotic marker and chromosomal morphology to identify anaphase and telophase GSCs, we found that the average ratios of old and new histone H3 (*n*=21), H4 (*n*=15), H2A (*n*=19), and the H3Rev (*n*=24) were nearly equal between the two sets of segregated sister chromatids in asymmetrically dividing GSCs (Fig 2A-D, quantified in Fig 2E). Given the intriguing similarity between GSCs and CBs in their old versus new H3 distribution patterns at prophase and prometaphase (Fig EV1B-C), we next examined old versus new H3 segregation patterns in anaphase and telophase CBs. It has been previously speculated that the daughter cell resulting from CB division that inherits more spectrosome, a germline-specific organelle (Koch & King, 1966, Lin & Spradling, 1995), eventually differentiates into the oocyte while the daughter cell that inherits less spectrosome commits to the nurse cell fate (de Cuevas & Spradling, 1998). By using both anti-α-spectrin and anti-1B1 to label the spectrosome structure, we distinguished the putative pre-oocyte versus pre-nurse cell between the two daughter cells derived from CB division. The average ratios of old and new histone H3 (*n*=19), H4 (*n*=24), H2A (*n*=23), and the H3Rev (*n*=20) were nearly equal between the two sets of segregated sister chromatids during CB division (Fig 2F-I, quantified in Fig 2J). Only a slight bias of H3 and H4 histone inheritance was detected in the putative future pre-nurse cell (less spectrosome, or spec-), compared to the possible future pre-oocyte cell (more spectrosome, or spec+, Figure 2J), but this was not statistically significant.

**Figure 2:**
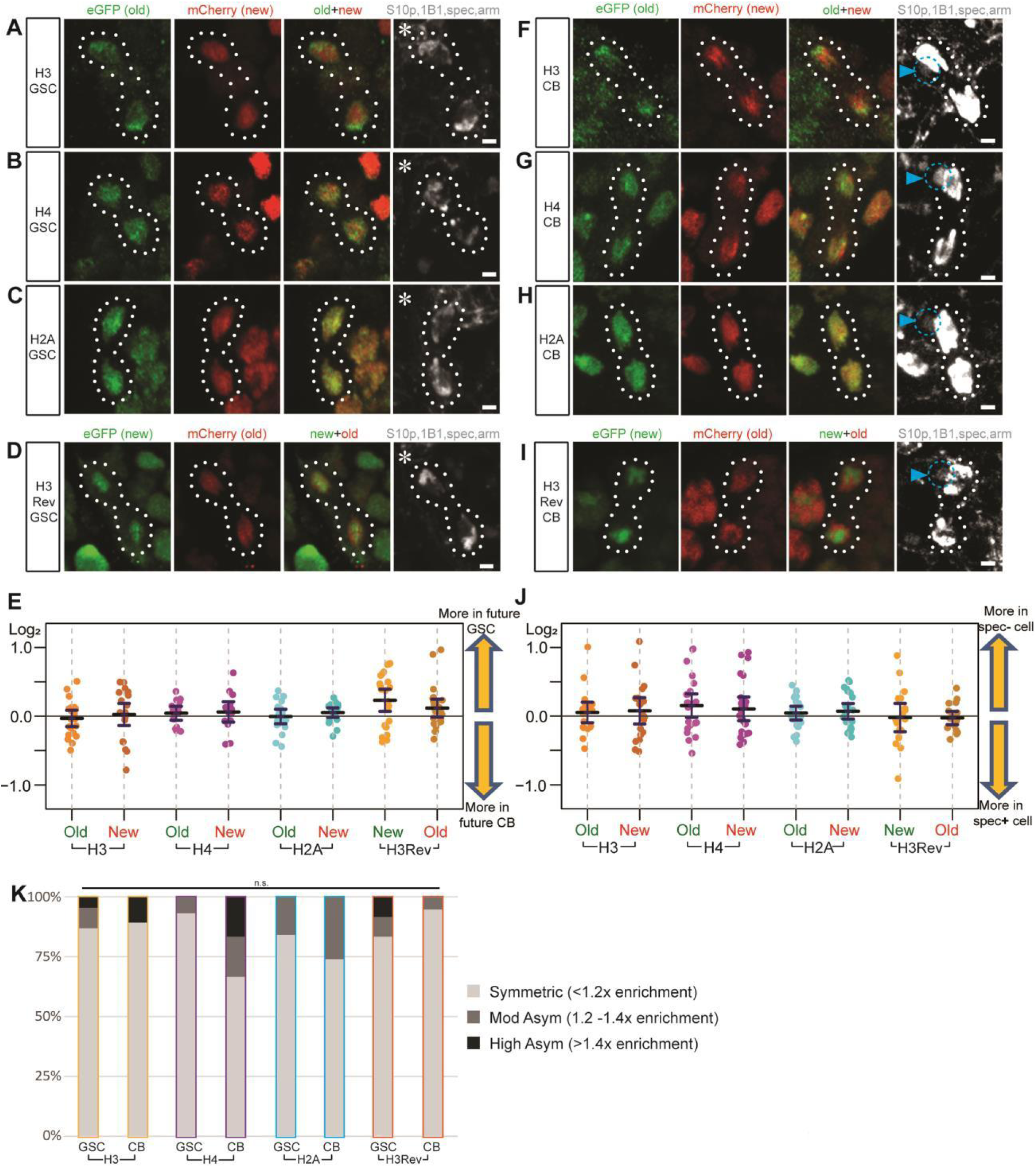
Mitotic GSCs and CBs exhibit non-overlapping old versus new histone H3 patterns, but overall symmetric old and new segregation patterns. (**A-C**) Telophase GSCs marked by H3S10ph (gray) expressing old (green) and new (red) histone H3 show non-overlapping patterns (**A**). Telophase GSCs expressing old (green) and new (red) histone H4 show moderate levels of overlap (**B**). GSCs expressing old (green) and new (red) histone H2A show high levels of overlap (**C**). (**D**) A control where the old and new fluorescent tags have been switched shows the non-overlapping patterns of old (red) versus new (green) H3Rev histone in mitotic telophase GSCs marked by H3S10ph (gray). (**E**) Quantification of log2 ratios of total old and new histone inherited by each future daughter cell of the GSC division, where a value of 0 is equal inheritance at exactly a 1:1 ratio. Values are mean ± 95% CI. (**F-H**) Telophase CBs marked by H3S10ph (gray) expressing old (green) and new (red) histone H3 also show non-overlapping patterns (**F**). Telophase CBs expressing old (green) and new (red) histone H4 (**G**) and H2A (**H**), which depict more overlap like GSCs. (**I**) A control with switched old and new fluorescent tags shows the non-overlapping old (red) versus new (green) H3Rev histone patterns in mitotic telophase CBs marked by H3S10p (gray). (**J**) Quantification of log2 ratios of total old and new histone inherited by each future daughter cell of the CB division, where a value of 0 is equal inheritance at exactly a 1:1 ratio. Values are mean ± 95% CI. (**K**) Summary of total old histone inherited in GSCs and CBs: <1.2-fold is symmetric, >1.2-fold but <1.4-fold is moderately asymmetric, and >1.4-fold is highly asymmetric. Asterisk: niche. Arrowheads (cyan): biased spec inheritance. Scale bars: 2μm. See relevant Supplemental Tables for individual data points for **Fig 2E** and **Fig 2J**.

Just like the separable old versus new H3 signals in prophase and prometaphase GSCs and CBs (Fig 1D, 1G, Fig EV1B-C), old and new H3 in anaphase and telophase GSCs and CBs (Fig 2A, 2D, 2F, 2I) still displayed non-overlapping patterns, compared to the overlapping old and new H2A patterns in anaphase and telophase GSCs and CBs (Fig 2C, 2H). Notably, old histones appeared to be more enriched at the poles in anaphase and telophase GSCs and CBs (Figure 2A, 2F), which are heterochromatin centromeric and pericentromeric regions. Recent studies have revealed an increased amount of old histone retention at the heterochromatin region when compared to that of the euchromatic region (Escobar et al., 2019). Additional studies will be needed to further explore whether there is any biological significance of this pattern in *Drosophila* female GSCs and CBs.

Interestingly, a subset of GSC and CB divisions results in two daughters with biased old versus new H3 and H4 inheritance, reflected by a wide distribution of H3 and H4 compared to H2A, which has a more clustered distribution (Fig 2E, 2J). For a more direct comparison, we classified all of the anaphase and telophase GSCs into three categories using previously established criteria for the degree of histone asymmetry (Ranjan et al., 2019, Xie et al., 2015).

The three categories include: (1) “symmetric,” where each daughter cell inherits a near equal amount of old histone (< 1.2-fold), (2) “moderately asymmetric” where one of the two daughter cells inherits between 1.2-1.4-fold old histone, and (3) “highly asymmetric” where one of the two daughter cells inherits more than 1.4-fold old histone (Fig 2K). Some H3 and H3Rev transgene-expressing GSCs as well as H4 transgene-expressing CBs showed more instances and a higher degree of asymmetric segregation pattern, whereas no H2A transgene-expressing GSC or CB showed such a pattern. H4 transgene-expressing CBs also showed more instances of asymmetric inheritance and a higher degree of asymmetry than H4 transgene-expressing GSCs. However, these differences were not statistically significant, so we concluded that during the GSC and CB divisions, histones are inherited in a globally symmetric manner.

Next, we performed similar analyses for old and new H3.3, with a shorter recovery time post-heat shock (see discussion above on recovery time, Fig EV2). Interestingly, old H3.3 showed significantly higher ratios of asymmetric inheritance patterns in GSC divisions (*n*=15) compared to the CB divisions (*n*=12, Fig EV2D). The sister chromatids segregated towards the GSC side inherited more of both old and new H3.3 than the sister chromatids segregated towards the CB side. This biased segregation pattern was not detected in the CB division (Fig EV2C), indicating that the future GSC has more H3.3 in total than the future CB during ACD of GSCs. Given the transcription-dependent incorporation mode of H3.3 (Ahmad & Henikoff, 2002b, Tagami et al., 2004), it is possible that this biased H3.3 inheritance contributes to or correlates with higher transcriptional activities in GSCs compared to CBs. These results are consistent with previous reports that stem cells maintain an overall more open chromatin state and their differentiated cells begin to “lock down” their chromatin towards a more restricted cell fate (Gaspar-Maia, Alajem et al., 2011, Golkaram, Jang et al., 2017, Turner, 2008, Yu, Wu et al., 2017).

### GSC-like cells from *bam* mutant ovaries recapitulate the non-overlapping old versus new H3 distribution patterns observed in wild-type female GSCs

In wild-type (WT) female ovaries, there are only 2-3 GSCs per ovariole and the M phase of GSCs only lasts for approximately 15 minutes in a ∼24-hour cell cycle (Ables & Drummond-Barbosa, 2013, Fuller & Spradling, 2007, Hinnant et al., 2017, Xie & Spradling, 2000). To obtain more GSC-like cells, we used *bag-of-marbles* (*bam*) mutant ovaries. Since the *bam* gene is necessary for female GSC differentiation, *bam* mutant ovaries are enriched with undifferentiated GSC-like cells [Fig 3A, (Chen & McKearin, 2003, Li, Minor et al., 2009, McKearin & Ohlstein, 1995, McKearin & Spradling, 1990)].

**Figure 3:**
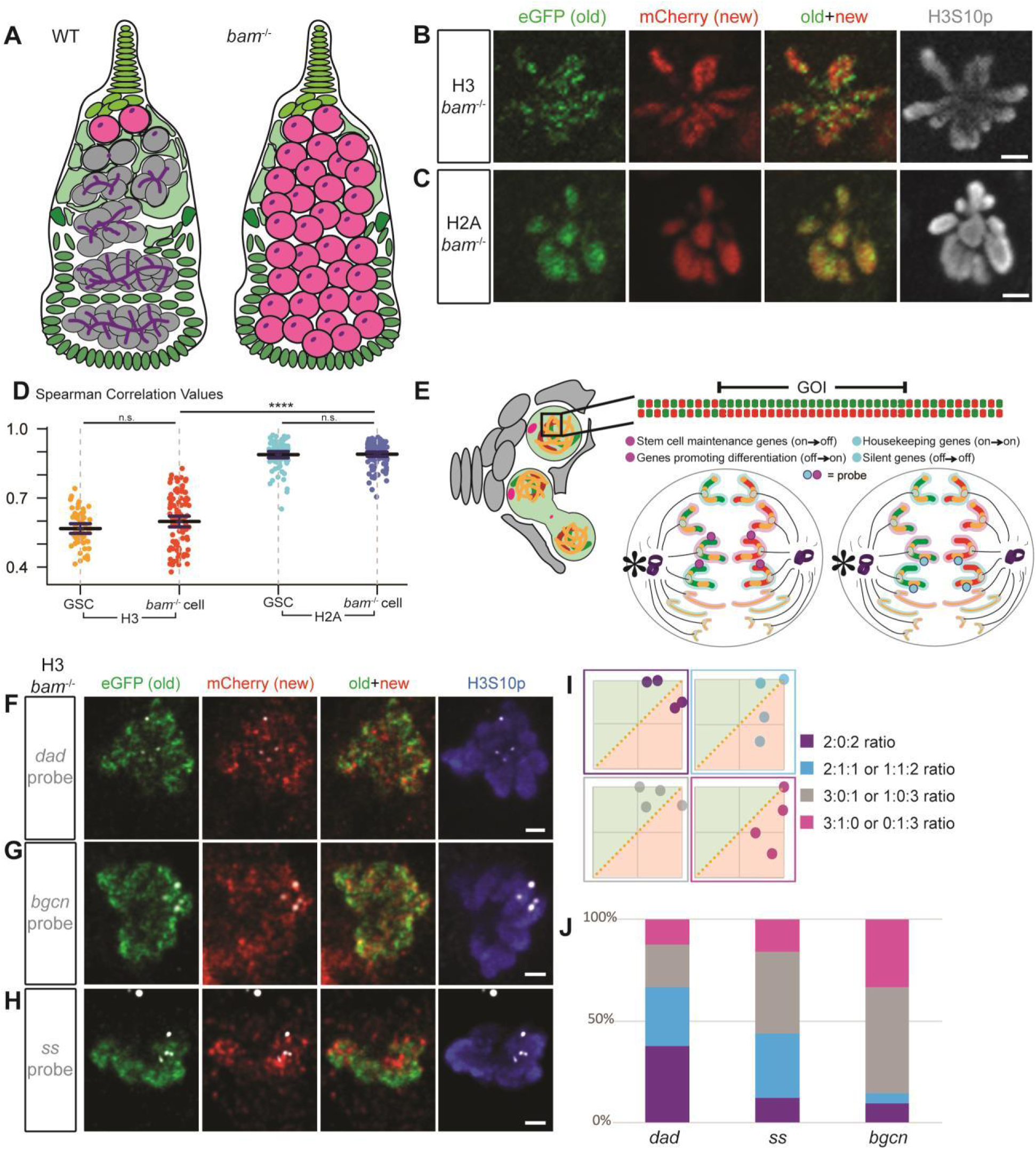
*bam* mutant germ cells recapitulate the separable old versus new H3 distribution in WT GSCs, and Oligopaint IF-FISH reveal distinct old versus new H3 at key genes. (**A**) A cartoon depicting a WT germarium with differentiating germline at different stages (gray) compared with a *bam* mutant ovariole filled with GSC-like cells (pink) without further differentiated germ cells. Undifferentiated cells can be identified by the presence of dotted spectrosome versus branched fusome structure (purple). (**B-C**) Non-overlapping patterns of old (green) versus new (red) histone H3 (**B**) and overlapping patterns for H2A (**C**) in mitotic prometaphase *bam* mutant cells marked by anti-H3S10ph (gray), similar to WT GSCs. Scale bars: 2μm. (**D**) Quantification of the overlap between old and new histones in late prophase and prometaphase *bam* mutant cells using Spearman correlation, also plotted with and compared to the WT GSC values from **Fig. 1L**. Values are mean ± 95% CI. *P*-value: pairwise ANOVA test with bonferroni correction. ****: *P*< 0.0001. n.s.: not significant. (**E**) A cartoon depicting the Oligopaint IF-FISH scheme to identify old versus new histone inheritance at single genomic loci in mitotic cells. (**Left**) A germarium containing a GSC in prometaphase, where the 3D chromatin structure may disguise local asymmetries at genomic regions of interest. (**Top**) A linearized gene of interest (GOI) displays local asymmetries. (**Bottom**) Two anaphase cells depict potential Oligopaint IF-FISH results. Maternal chromosomes are outlined in pink and paternal ones in blue. (**Bottom Left**) Probes recognize genes (magenta) that change their epigenetic state in a 2:2 ratio with biased old:new H3-enriched regions. In this instance, each duplicated sister chromatid from either maternal or paternal chromosome has an “agreement” on old (green, inherited by GSC indicated by an asterisk) versus new (red, inherited by the CB) H3. (**Bottom Right**) Probes recognize genes (cyan) that maintain their epigenetic state associated with more symmetric regions (yellow) that do not have a strong bias for either old or new H3. (**F-H**) Examples showing old (green) versus new (red) H3 distribution at a single genomic locus labeled with fluorescent probes (gray) for *dad* gene (**F**), *bgcn* gene (**G**), and *ss* gene (**H**) in *bam* mutant GSC-like cells at prometaphase marked by anti-H3S10ph (blue). Scale bars: 1μm. (**I**) Examples of scatter plots showing probe’s association with normalized old H3 (green, above X=Y line) versus new H3 (red, below X=Y line) and the (i):(ii):(iii) ratios that result from them, where (i) is the number of FISH signals more associated with old histone, (ii) is the number of FISH signals that had equal association with both old and new histone, and (iii) is the number of FISH signals more associated with new histone. (**J**) Quantification of the probe association ratios with old versus new H3 for each candidate gene in *bam* mutant germ cells. See relevant Supplemental Tables for individual data points for **Fig 3D** and **Fig 3J**.

Next, we performed similar experiments using the dual-color histone H3 and H2A transgenes in *bam* mutant ovaries [*bam*^Δ86^*/bam*^1^, (Bopp, Horabin et al., 1993, McKearin & Spradling, 1990)] and compared the results with those from WT GSCs. Similar to WT GSCs, GSC-like cells in *bam* mutant ovaries at late prophase to prometaphase displayed non-overlapping old versus new H3 patterns (Fig 3B), but largely overlapping old versus new H2A patterns (Fig 3C). Quantification using Spearman’s rank correlation coefficient showed an average of 0.598 for H3 (*n*= 98) and 0.890 for H2A (*n*= 116). Both coefficient values for H3 and H2A in *bam* mutant germ cells were indistinguishable from those in WT GSCs (Fig 3D).

Furthermore, *bam* mutant germ cells showed significantly different correlation coefficients between old versus new H3 and old versus new H2A, similar to WT GSCs (Fig 3D). Together, these results suggest that the non-overlapping old versus new H3 distribution in WT female GSCs are recapitulated in *bam* mutant germ cells, which greatly facilitate higher throughput image acquisition and data analyses. Furthermore, the similarity of old versus new histone distribution patterns among WT GSCs, WT CBs, and *bam* mutant germ cells (data re-plotted in Fig EV3D for direct comparison) shed light on a long-held debate in the field whether *bam* mutant germ cells resemble more like GSCs or CBs (Casanueva & Ferguson, 2004, McKearin & Spradling, 1990, Shen, Weng et al., 2009). Here, our results show that WT GSCs, WT CBs, and *bam* mutant germ cells have more similarities than differences, and that the non-overlapping old versus new H3 distribution patterns gradually diminish along with germline differentiation after the CB stage (Fig EV1B-C). In accordance with this observation, it has been shown that other cellular features, such as cell cycle timing, also change gradually during female germline differentiation (Hinnant et al., 2017). Collectively, these observations confirm a step-wise cellular differentiation pathway in the female GSC lineage.

### Oligopaint IF-FISH reveals differential old versus new H3 distribution at key genes for maintaining stem cell fate but not at constitutively active or silent genes

The non-overlapping old versus new H3 distribution patterns in WT GSCs and *bam* mutant germ cells introduce the intriguing possibility that old and new H3 could be differentially distributed at specific genomic loci, rather than globally across the entire genome. We hypothesize that old and new H3 are differentially distributed and inherited at genes that are important for maintaining the stem cell state (i.e. stemness genes) or for promoting cellular differentiation (i.e. differentiation genes). For example, the self-renewed stem cell could inherit old histones with post-translational modifications to maintain the active expression of the stemness genes, while the differentiating daughter cell could carry new histones without these modifications (or with different modifications) so that the expression of stemness genes are silenced. Other genes such as constitutively active genes or non-germline lineage genes might display a more symmetric histone inheritance pattern, which would allow both daughter cells to achieve a similar gene expression pattern after ACD.

To visualize old versus new histones at specific genomic loci and between sister chromatids, we combined the dual color histone-labeling scheme with DNA Oligopaint FISH technology (Beliveau, Joyce et al., 2012, Joyce, Apostolopoulos et al., 2013) (Fig 3E). Our candidate target genes included the stemness gene *daughters against dpp* (*dad*) (Casanueva & Ferguson, 2004, Kirilly et al., 2011, Xie & Spradling, 1998), the germline differentiation gene *bam* (McKearin & Ohlstein, 1995, McKearin & Spradling, 1990), an actively transcribed gene in both GSCs and CBs called *benign gonial cell neoplasm* [*bgcn*, Fig EV3A-B, (Gönczy, Matunis et al., 1997, Lavoie, Ohlstein et al., 1999)], and a neuronal gene that is completely silenced in the entire female germline lineage, *spineless* (*ss*) (Gan, Chepelev et al., 2010, Struhl, 1982, Wernet, Mazzoni et al., 2006). Among these candidates, *dad, bam*, and *ss* are located on the right arm of the 3^rd^ chromosome (3R), and *bgcn* is located on the right arm of the 2^nd^ chromosome (2R), all of which are autosomes in fly cells.

Since the non-overlapping old versus new H3 distribution pattern in *bam* mutant germ cells resembled that in WT GSCs (Fig 3B), we first used *bam* mutant ovaries to achieve a higher number of cells where histone distribution patterns at candidate genomic regions could be studied. We imaged mitotic cells using high-resolution Airyscan microscopy (Sivaguru, Urban et al., 2018) to increase the spatial resolution and resolved the candidate genomic regions in prophase cells into four fluorescence *in situ* hybridization (FISH) “dots”, two of each for the replicated maternal and paternal chromosomes, respectively. This allowed us to examine how the candidate gene loci were associated with old versus new histone H3. We hypothesize that if histones are differentially distributed at stemness and differentiation genes, we would detect two FISH signals associated with old H3 and two FISH signals associated with new H3. However, if the histones are symmetrically inherited, we would not be able to detect this 2:2 ratio of old:new H3 association at candidate gene loci.

Using Oligopaint IF-FISH, we first examined each candidate gene locus in *bam* mutant germ cells at late prophase when the four distinct FISH dots were most likely detectable. The mitotic chromosomes were co-immunostained with antibodies against H3S10ph, the tag for old H3, and the tag for new H3. As shown previously (Fig 3B), old (GFP) and new (mCherry) H3 displayed non-overlapping patterns in mitotic *bam* mutant germ cells (Fig 3F-H: old + new signals), and the four FISH signals indicate duplicated sister chromatids for both maternal and paternal chromosomes. It is worth noting that homologous chromosomes are not paired in the early-stage female germline, unlike somatic cells in *Drosophila* (Joyce et al., 2013). We then asked whether these four FISH signals show any distinct patterns associated with old versus new H3. We reasoned that if two dots displayed an association with old histone and the other two were enriched with new histone, there were two possibilities: (1) One sister chromatid from the maternal chromosome and one sister chromatid from the paternal chromosome are associated with old H3, while the others are associated with new H3 (left panel in Figure 3E); (2) Both sister chromatids from either the maternal or paternal chromosome are associated with old H3, and both of the remaining paternal or maternal chromatids are associated with new H3 (Fig EV3C).

To examine whether the FISH signals were more associated with old (green) or new (red) histones, we measured the mean intensity values for each channel (see Materials and Methods). We then normalized both the red and green channel values by making each highest mean intensity value equal to 1 and dividing the other three signal values accordingly to create percentages for each one. Next, we plotted the normalized mean intensity values for each probe signal in a two-dimensional (2D) scatter plot with the X-axis representing the new H3 signal and the Y-axis representing the old H3 signal. Finally, we drew a diagonal line at X=Y and determined whether each probe signal was more associated with old H3 or with new H3 (Fig. 3I, Materials and Methods), followed by classifying all scatterplot patterns with (i):(ii):(iii) ratios, where (i) is the number of FISH signals more associated with the old histone (above the X=Y line), (ii) is the number of FISH signals that with an equal association between old and new histone (on the X=Y line), and (iii) is the number of FISH signals more associated with new histone (below the X=Y line). For the stemness *dad* gene (Fig 3F), ∼38% of the *bam* germ cells showed two dots associated with the old histone and two with the new histone (a 2:0:2 ratio, *n* = 24, quantified in Fig 3J). By contrast, when probing for *bgcn* (Fig 3G, *n*= 21) and *ss* (Fig 3H, *n*= 25), fewer cells (9.5% and 12%, respectively) displayed this 2:0:2 ratio (Fig 3J). These results suggest that the 2:0:2 histone association pattern is more frequently detected in genes that must change expression between the two daughter cells, such as *dad*. For genes that do not change expression between the two daughter cells, such as *ss* and *bgcn*, this ratio is less frequent (Fig 3J). Notably, both *bgcn* and *ss* share similar expression profiles between GSC and CB, although *bgcn* is actively expressed (on) and *ss* is repressed (off).

A caveat in *bam* mutant cells is that the genetic lesions at the *bam* genomic locus might complicate FISH probing at the *bam* gene region. Moreover, in *bam* mutant ovaries, no ACD occurs because all of the germ cells overproliferate as GSC-like cells without differentiation. Therefore, to study histone inheritance patterns during stem cell ACD, we examined Oligopaint IF-FISH signals with old versus new H3 segregation patterns in asymmetrically dividing WT GSCs. Since for both the constitutively active *bgcn* gene and the silent *ss* gene, very few GSC-like cells displayed the 2:0:2 ratio, we next focused on differentially expressed *dad* gene and *bam* gene in WT GSCs.

### Oligopaint IF-FISH reveals differential old versus new H3 inheritance at key genes for maintaining stem cell fate or for promoting differentiation in WT GSCs

In mitotic WT GSCs, we detected non-overlapping old versus new H3 distribution patterns (Fig 1D, 1G, 1L, Fig 2A, 2D). Here, we also visualized the localization of old versus new histone signals at candidate genes using Oligopaint IF-FISH. Notably, FISH signals from duplicated sister chromatids of the maternal and paternal chromosomes often result in two adjacent dots (an example of a prophase GSC shown in Fig 4A, and examples of telophase GSCs shown in Fig 4C-D). In prophase GSCs, this is likely due to the action of cohesin proteins between sister chromatids (Brooker & Berkowitz, 2014). In anaphase and telophase GSCs, this is probably a result of the co-segregation of maternal and paternal chromosomes via the pulling force of the mitotic spindle. Co-segregation of maternal and paternal autosomes has been formerly reported in *Drosophila* male GSCs (Yadlapalli & Yamashita, 2013).

**Figure 4:**
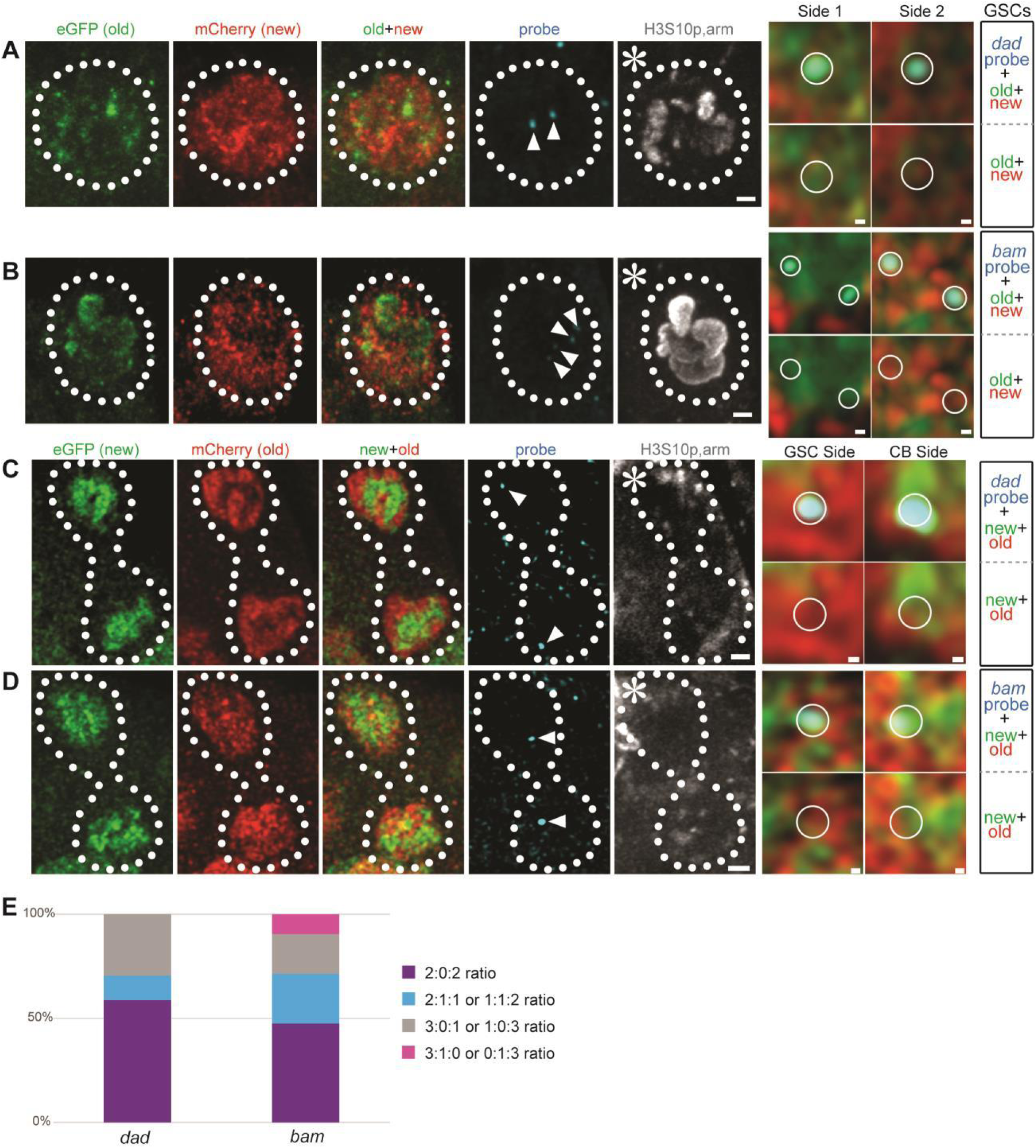
Oligopaint IF-FISH reveals differential old versus new H3 inheritance at key genes for maintaining stem cell fate or promoting differentiation in WT female GSCs. (**A-B**) Old (green) versus new (red) histone patterns with a single genomic locus labeled with fluorescent probes (cyan) for *dad* (**A**) and *bam* (**B**) in WT prometaphase female GSCs, marked by anti-H3S10ph (gray). (**C-D**) Old (green) versus new (red) histone inheritance patterns with single genes labeled with fluorescent probes (cyan) for *dad* (**C**) and *bam* (**D**) in WT telophase female GSCs, marked by anti-H3S10ph (gray). (**E**) Quantification of the probe association ratios with old versus new H3 for each candidate gene in WT female GSCs. Asterisk: niche. Arrowheads: probe signal. Scale bars: 1μm. Zoomed-in images of each probe are displayed to the right of each figure panel. Scale bars: 0.1μm. See relevant Supplemental Table for individual data points for **Fig 4E**.

Interestingly, when all four dots of the stemness *dad* gene are resolvable, ∼59% of the WT mitotic GSCs showed two dots associated with the old histone and two dots with the new histone (Fig 4A, *n*= 17, quantified in Fig 4E). For the germline differentiation *bam* gene, ∼48% of WT mitotic GSCs showed two dots associated with the old histone and two with the new histone (Fig 4B, *n*= 21, quantified in Fig 4E). However, the normalization scheme could potentially create a situation in which one dot has both the highest GFP and the highest mCherry signal, leading to a 2:1:1 pattern (see Materials and Methods), which is likely due to the high condensation of chromosomes in mitotic cells. If we consider both the 2:0:2 and the 2:1:1 pattern, approximately 71% of *dad* and 71% of *bam* FISH signals have a preferential association with old versus new histones. For the ∼30% of signals where a preferential association was not detected with either old or new histones, it is possible that fewer labeled histones were incorporated at that genomic region, or that the chromatin was folded and condensed in such a manner that immunostaining was unable to detect it.

Finally, for differentially expressed *dad* and *bam* genes, we examined the segregation patterns of old versus new histone H3 at their corresponding genomic loci in anaphase or telophase WT GSCs, where sister chromatids can be easily distinguished as they segregate (Fig 4C for the *dad* gene locus and Fig 4D for the *bam* gene locus). When probing at the *dad* locus, the future GSC side had predominantly old H3 while the CB side mainly inherited new histones at this gene locus ∼75% of the time (Fig 4C, *n*= 12). A similar pattern was observed in ∼67% GSCs at the *bam* locus (Fig 4D, *n*= 12). Specifically, out of the 10 telophase cells displaying a 2:0:2 ratio at *dad* locus (*n*= 12), 90% showed old H3 on GSC side and new H3 on CB side. Additionally, out of the 10 telophase cells displaying a 2:0:2 ratio at *bam* locus (*n*= 12), 80% showed old H3 on GSC side and new H3 on CB side. Collectively, these data indicate a differential association of old versus new histone at specific gene loci, followed by the preferential segregation of sister chromatids with old H3-enriched sister chromatids segregating toward the GSC side and new H3-enriched sister chromatids segregating toward the CB side.

In summary, in this study we detected differentially distributed old versus new histone H3 in prophase and prometaphase early-stage female germ cells. This differential distribution has molecular specificity that is detectable in H3 but not in H2A, as well as cellular specificity for GSCs and CBs but not for further differentiated CCs. Using Oligopaint IF-FISH, this differential distribution is likely associated with the key genes either for maintaining the stem cell state or for promoting differentiation, which could establish distinct epigenomes in the two daughter cells arising from ACD. The combination of using differential labeling for old versus new histone with Oligopaint IF-FISH is a technique that can be adapted and applied to many other systems, including induced asymmetrically dividing mouse embryonic stem cells where differential histone H3 distribution patterns is also detected (Ma, Trieu et al., 2020). Application of this technique to other systems will provide valuable insight into how ACD regulates cell fate decisions in multicellular organisms, which likely involves changing the expression of a handful of key genes, while maintaining similar expression patterns for other genes required for homeostasis and lineage specificity.

Previous studies have shown that recycled old histones can be biasedly deposited to either the leading or lagging strands following DNA replication under different conditions (Petryk et al., 2018, Roufa & Marchionni, 1982, Seale, 1976, Seidman, Levine et al., 1979, Wooten et al., 2019, Yu et al., 2018). While the origins of DNA replication have yet to be determined in the majority of multicellular organism cell types, it is well known that transcription can directly affect the localization of pre-replication complexes (Cayrou, Coulombe et al., 2011, Vashee, Cvetic et al., 2003). This introduces the possibility that transcription may affect the location of replication origins, the length of the replicons, and biased old versus new histone incorporation at the corresponding genomic loci (Kahney, Ranjan et al., 2017), in a cell type-specific manner. Future studies using molecular genetics, genomic, cell biology and biophysical tools will be needed to directly test this speculation in different cell types and to address this fundamental question in developmental biology.

## Acknowledgements

We thank Drs. M. Van Doren, J. Gall, D. Drummond-Barbosa, A. Spradling, and X.C. lab members for insightful suggestions. We also thank T. Wu (Harvard) and E. Joyce (UPenn) and their lab members for thoughtful discussions and reagents. We thank Johns Hopkins Integrated Imaging Center for confocal imaging. This work was supported by F31 GM122339 (E.W.K.), F31EY026786 (K.V.L.), R01EY025598 (R.J.), R01GM112008 and R35GM127075, the Howard Hughes Medical Institute, the David and Lucile Packard Foundation, and Johns Hopkins University startup funds (X.C.).

## Author contribution

E.W.K. and X.C. conceptualized the study; K.V.L. and R.J. provided Oligopaint training and some relevant reagents; E.W.K. and L.S. performed all the experiments and data analyses; E.W.K. and X.C. wrote the manuscript.

## Competing interests

The authors declare no competing interests.

## REFERENCES

Ables ET, Drummond-Barbosa D (2013) Cyclin E controls Drosophila female germline stem cell maintenance independently of its role in proliferation by modulating responsiveness to niche signals. Development 140: 530–40

Ahmad K, Henikoff S (2002a) Histone H3 variants specify modes of chromatin assembly. Proc Natl Acad Sci U S A 99 Suppl 4: 16477–84

Ahmad K, Henikoff S (2002b) The histone variant H3.3 marks active chromatin by replication-independent nucleosome assembly. Mol Cell 9: 1191–200

Ahmad K, Henikoff S (2018) No strand left behind. Science 361: 1311–1312

Alabert C, Barth TK, Reverón-Gómez N, Sidoli S, Schmidt A, Jensen ON, Imhof A, Groth A (2015) Two distinct modes for propagation of histone PTMs across the cell cycle. Genes Dev 29: 585–90

Alabert C, Groth A (2012) Chromatin replication and epigenome maintenance. Nat Rev Mol Cell Biol 13: 153–67

Allis CD, Jenuwein T (2016) The molecular hallmarks of epigenetic control. Nat Rev Genet 17: 487–500

Baylin SB, Ohm JE (2006) Epigenetic gene silencing in cancer - a mechanism for early oncogenic pathway addiction? Nat Rev Cancer 6: 107–16

Beliveau BJ, Joyce EF, Apostolopoulos N, Yilmaz F, Fonseka CY, McCole RB, Chang Y, Li JB, Senaratne TN, Williams BR, Rouillard JM, Wu CT (2012) Versatile design and synthesis platform for visualizing genomes with Oligopaint FISH probes. Proc Natl Acad Sci U S A 109: 21301–6

Bopp D, Horabin JI, Lersch RA, Cline TW, Schedl P (1993) Expression of the Sex-lethal gene is controlled at multiple levels during Drosophila oogenesis. Development 118: 797–812

Brooker AS, Berkowitz KM (2014) The roles of cohesins in mitosis, meiosis, and human health and disease. Methods Mol Biol 1170: 229–66

Casanueva MO, Ferguson EL (2004) Germline stem cell number in the Drosophila ovary is regulated by redundant mechanisms that control Dpp signaling. Development 131: 1881–90

Cayrou C, Coulombe P, Vigneron A, Stanojcic S, Ganier O, Peiffer I, Rivals E, Puy A, Laurent-Chabalier S, Desprat R, Mechali M (2011) Genome-scale analysis of metazoan replication origins reveals their organization in specific but flexible sites defined by conserved features. Genome Res 21: 1438–49

Chambers SM, Shaw CA, Gatza C, Fisk CJ, Donehower LA, Goodell MA (2007) Aging hematopoietic stem cells decline in function and exhibit epigenetic dysregulation. PLoS Biol 5: e201

Chen D, McKearin DM (2003) A discrete transcriptional silencer in the bam gene determines asymmetric division of the Drosophila germline stem cell. Development 130: 1159–70

Dattoli AA, Carty, B.L., Kochendoerfer, A.M.,, Morgan, C., Walshe, A.E. and Dunleavy, E.M. (2020) Asymmetric assembly of centromeres epigenetically regulates stem cell fate. J Cell Biol 219: e201910084

de Cuevas M, Spradling AC (1998) Morphogenesis of the Drosophila fusome and its implications for oocyte specification. Development 125: 2781–9

Escobar TM, Oksuz O, Saldana-Meyer R, Descostes N, Bonasio R, Reinberg D (2019) Active and Repressed Chromatin Domains Exhibit Distinct Nucleosome Segregation during DNA Replication. Cell 179: 953–963 e11

Feinberg AP, Ohlsson R, Henikoff S (2006) The epigenetic progenitor origin of human cancer. Nat Rev Genet 7: 21–33

Fitzsimons CP, van Bodegraven E, Schouten M, Lardenoije R, Kompotis K, Kenis G, van den Hurk M, Boks MP, Biojone C, Joca S, Steinbusch HW, Lunnon K, Mastroeni DF, Mill J, Lucassen PJ, Coleman PD, van den Hove DL, Rutten BP (2014) Epigenetic regulation of adult neural stem cells: implications for Alzheimer’s disease. Mol Neurodegener 9: 25

Fredly H, Gjertsen BT, Bruserud O (2013) Histone deacetylase inhibition in the treatment of acute myeloid leukemia: the effects of valproic acid on leukemic cells, and the clinical and experimental evidence for combining valproic acid with other antileukemic agents. Clin Epigenetics 5: 12

Fuller MT, Spradling AC (2007) Male and female Drosophila germline stem cells: two versions of immortality. Science 316: 402–4

Gan H, Serra-Cardona A, Hua X, Zhou H, Labib K, Yu C, Zhang Z (2018) The Mcm2-Ctf4-Polalpha Axis Facilitates Parental Histone H3-H4 Transfer to Lagging Strands. Mol Cell 72: 140–151 e3

Gan Q, Chepelev I, Wei G, Tarayrah L, Cui K, Zhao K, Chen X (2010) Dynamic regulation of alternative splicing and chromatin structure in Drosophila gonads revealed by RNA-seq. Cell Res 20: 763–83

Gaspar-Maia A, Alajem A, Meshorer E, Ramalho-Santos M (2011) Open chromatin in pluripotency and reprogramming. Nat Rev Mol Cell Biol 12: 36–47

Golkaram M, Jang J, Hellander S, Kosik KS, Petzold LR (2017) The Role of Chromatin Density in Cell Population Heterogeneity during Stem Cell Differentiation. Sci Rep 7: 13307

Gönczy P, Matunis E, DiNardo S (1997) bag-of-marbles and benign gonial cell neoplasm act in the germline to restrict proliferation during Drosophila spermatogenesis. Development 124: 4361–71

Hardy RW, Tokuyasu KT, Lindsley DL (1981) Analysis of spermatogenesis in Drosophila melanogaster bearing deletions for Y-chromosome fertility genes. Chromosoma 83: 593–617

Hardy RW, Tokuyasu KT, Lindsley DL, Garavito M (1979) The germinal proliferation center in the testis of Drosophila melanogaster. J Ultrastruct Res 69: 180–90

Hinnant TD, Alvarez AA, Ables ET (2017) Temporal remodeling of the cell cycle accompanies differentiation in the Drosophila germline. Dev Biol 429: 118–131

Holtzman S, Miller D, Eisman R, Kuwayama H, Niimi T, Kaufman T (2010) Transgenic tools for members of the genus Drosophila with sequenced genomes. Fly (Austin) 4: 349–62

Hsu HJ, LaFever L, Drummond-Barbosa D (2008) Diet controls normal and tumorous germline stem cells via insulin-dependent and -independent mechanisms in Drosophila. Dev Biol 313: 700–12

Huang H, Sabari BR, Garcia BA, Allis CD, Zhao Y (2014) SnapShot: histone modifications. Cell 159: 458–458 e1

Jacobs JJ, van Lohuizen M (2002) Polycomb repression: from cellular memory to cellular proliferation and cancer. Biochim Biophys Acta 1602: 151–61

Janssen KA, Sidoli S, Garcia BA (2017) Recent Achievements in Characterizing the Histone Code and Approaches to Integrating Epigenomics and Systems Biology. Methods Enzymol 586: 359–378

Jin B, Li Y, Robertson KD (2011) DNA methylation: superior or subordinate in the epigenetic hierarchy? Genes Cancer 2: 607–17

Joyce EF, Apostolopoulos N, Beliveau BJ, Wu CT (2013) Germline progenitors escape the widespread phenomenon of homolog pairing during Drosophila development. PLoS Genet 9: e1004013

Kahney EW, Ranjan R, Gleason RJ, Chen X (2017) Symmetry from Asymmetry or Asymmetry from Symmetry? Cold Spring Harb Symp Quant Biol 82: 305–318

King RC (1957) Oogenesis in adult Drosophila melanogaster. II. Stage distribution as a function of age. Growth 21: 95–102

Kirilly D, Wang S, Xie T (2011) Self-maintained escort cells form a germline stem cell differentiation niche. Development 138: 5087–97

Koch EA, King RC (1966) The origin and early differentiation of the egg chamber of Drosophila melanogaster. J Morphol 119: 283–303

Kouzarides T (2007) Chromatin modifications and their function. Cell 128: 693–705

Lavoie CA, Ohlstein B, McKearin DM (1999) Localization and function of Bam protein require the benign gonial cell neoplasm gene product. Dev Biol 212: 405–13

Lee JT (2012) Epigenetic regulation by long noncoding RNAs. Science 338: 1435–9

Li Y, Minor NT, Park JK, McKearin DM, Maines JZ (2009) Bam and Bgcn antagonize Nanos-dependent germ-line stem cell maintenance. Proc Natl Acad Sci U S A 106: 9304–9

Lin H, Spradling AC (1995) Fusome asymmetry and oocyte determination in Drosophila. Dev Genet 16: 6–12

Lin S, Yuan ZF, Han Y, Marchione DM, Garcia BA (2016) Preferential Phosphorylation on Old Histones during Early Mitosis in Human Cells. J Biol Chem 291: 15342–57

Lyko F, Ramsahoye BH, Jaenisch R (2000) DNA methylation in Drosophila melanogaster. Nature 408: 538–40

Ma B, Trieu TJ, Cheng J, Zhou S, Tang Q, Xie J, Liu JL, Zhao K, Habib SJ, Chen X (2020) Differential Histone Distribution Patterns in Induced Asymmetrically Dividing Mouse Embryonic Stem Cells. Cell Rep 32: 108003

McKearin D, Ohlstein B (1995) A role for the Drosophila bag-of-marbles protein in the differentiation of cystoblasts from germline stem cells. Development 121: 2937–47

McKearin DM, Spradling AC (1990) bag-of-marbles: a Drosophila gene required to initiate both male and female gametogenesis. Genes Dev 4: 2242–51

Petryk N, Dalby M, Wenger A, Stromme CB, Strandsby A, Andersson R, Groth A (2018) MCM2 promotes symmetric inheritance of modified histones during DNA replication. Science 361: 1389–1392

Probst AV, Dunleavy E, Almouzni G (2009) Epigenetic inheritance during the cell cycle. Nat Rev Mol Cell Biol 10: 192–206

Ranjan R, Snedeker J, Chen X (2019) Asymmetric Centromeres Differentially Coordinate with Mitotic Machinery to Ensure Biased Sister Chromatid Segregation in Germline Stem Cells. Cell Stem Cell 25: 666–681 e5

Reveron-Gomez N, Gonzalez-Aguilera C, Stewart-Morgan KR, Petryk N, Flury V, Graziano S, Johansen JV, Jakobsen JS, Alabert C, Groth A (2018) Accurate Recycling of Parental Histones Reproduces the Histone Modification Landscape during DNA Replication. Mol Cell 72: 239–249 e5

Ringrose L, Paro R (2004) Epigenetic regulation of cellular memory by the Polycomb and Trithorax group proteins. Annu Rev Genet 38: 413–43

Roufa DJ, Marchionni MA (1982) Nucleosome segregation at a defined mammalian chromosomal site. Proc Natl Acad Sci U S A 79: 1810–4

Schober P, Boer C, Schwarte LA (2018) Correlation Coefficients: Appropriate Use and Interpretation. Anesth Analg 126: 1763–1768

Seale RL (1976) Studies on the mode of segregation of histone nu bodies during replication in HeLa cells. Cell 9: 423–9

Seidman MM, Levine AJ, Weintraub H (1979) The asymmetric segregation of parental nucleosomes during chrosome replication. Cell 18: 439–49

Serra-Cardona A, Zhang Z (2018) Replication-Coupled Nucleosome Assembly in the Passage of Epigenetic Information and Cell Identity. Trends Biochem Sci 43: 136–148

Shen R, Weng C, Yu J, Xie T (2009) eIF4A controls germline stem cell self-renewal by directly inhibiting BAM function in the Drosophila ovary. Proc Natl Acad Sci U S A 106: 11623–8

Sivaguru M, Urban MA, Fried G, Wesseln CJ, Mander L, Punyasena SW (2018) Comparative performance of airyscan and structured illumination superresolution microscopy in the study of the surface texture and 3D shape of pollen. Microsc Res Tech 81: 101–114

Smith AV, Orr-Weaver TL (1991) The regulation of the cell cycle during Drosophila embryogenesis: the transition to polyteny. Development 112: 997–1008

Struhl G (1982) Spineless-aristapedia: a homeotic gene that does not control the development of specific compartments in Drosophila. Genetics 102: 737–49

Tagami H, Ray-Gallet D, Almouzni G, Nakatani Y (2004) Histone H3.1 and H3.3 complexes mediate nucleosome assembly pathways dependent or independent of DNA synthesis. Cell 116: 51–61

Tokuyasu KT, Peacock WJ, Hardy RW (1977) Dynamics of spermiogenesis in Drosophila melanogaster. VII. Effects of segregation distorter (SD) chromosome. J Ultrastruct Res 58: 96–107

Tran V, Lim C, Xie J, Chen X (2012) Asymmetric division of Drosophila male germline stem cell shows asymmetric histone distribution. Science 338: 679–82

Turner BM (2002) Cellular memory and the histone code. Cell 111: 285–91

Turner BM (2008) Open chromatin and hypertranscription in embryonic stem cells. Cell Stem Cell 2: 408–10

Vashee S, Cvetic C, Lu W, Simancek P, Kelly TJ, Walter JC (2003) Sequence-independent DNA binding and replication initiation by the human origin recognition complex. Genes Dev 17: 1894–908

Wernet MF, Mazzoni EO, Çelik A, Duncan DM, Duncan I, Desplan C (2006) Stochastic spineless expression creates the retinal mosaic for colour vision. Nature 440: 174–80

Wooten M, Snedeker J, Nizami ZF, Yang X, Ranjan R, Urban E, Kim JM, Gall J, Xiao J, Chen X (2019) Asymmetric histone inheritance via strand-specific incorporation and biased replication fork movement. Nat Struct Mol Biol 26: 732–743

Xie J, Wooten M, Tran V, Chen BC, Pozmanter C, Simbolon C, Betzig E, Chen X (2015) Histone H3 Threonine Phosphorylation Regulates Asymmetric Histone Inheritance in the Drosophila Male Germline. Cell 163: 920–33

Xie T, Spradling AC (1998) decapentaplegic is essential for the maintenance and division of germline stem cells in the Drosophila ovary. Cell 94: 251–60

Xie T, Spradling AC (2000) A niche maintaining germ line stem cells in the Drosophila ovary. Science 290: 328–30

Xu M, Wang W, Chen S, Zhu B (2011) A model for mitotic inheritance of histone lysine methylation. EMBO Rep 13: 60–7

Yadlapalli S, Yamashita YM (2013) Chromosome-specific nonrandom sister chromatid segregation during stem-cell division. Nature 498: 251–4

Young NL, DiMaggio PA, Garcia BA (2010) The significance, development and progress of high-throughput combinatorial histone code analysis. Cellular and Molecular Life Sciences 67: 3983–4000

Yu C, Gan H, Serra-Cardona A, Zhang L, Gan S, Sharma S, Johansson E, Chabes A, Xu RM, Zhang Z (2018) A mechanism for preventing asymmetric histone segregation onto replicating DNA strands. Science 361: 1386–1389

Yu X, Wu C, Bhavanasi D, Wang H, Gregory BD, Huang J (2017) Chromatin dynamics during the differentiation of long-term hematopoietic stem cells to multipotent progenitors. Blood Adv 1: 887–898

Zee BM, Britton LM, Wolle D, Haberman DM, Garcia BA (2012) Origins and formation of histone methylation across the human cell cycle. Mol Cell Biol 32: 2503–14

Zhang G, Huang H, Liu D, Cheng Y, Liu X, Zhang W, Yin R, Zhang D, Zhang P, Liu J, Li C, Liu B, Luo Y, Zhu Y, Zhang N, He S, He C, Wang H, Chen D (2015) N6-methyladenine DNA modification in Drosophila. Cell 161: 893–906

